# Rapid and precise quantification of lymphocyte iron content by single cell inductively coupled plasma mass spectrometry

**DOI:** 10.1101/2024.11.11.623006

**Authors:** Philip Holdship, Megan R. Teh, Michalina Mazurczyk, Huei-Wen Chuang, Giulia Pironaci, Robert G. Hilton, David Price, Jon Wade, Hal Drakesmith

## Abstract

Metals facilitate catalysis during cellular metabolism, but heterogeneity of metal content at single-cell level within and between cell populations is poorly characterized. This is important because deficiencies of biometals, for example iron, are enormously prevalent worldwide. Here we quantify metal content of single-cells using inductively-coupled plasma mass spectrometry. To develop the method, we used rhodium and iridium-intercalated Jurkat cells, obtaining >0.96% r^2^ cross-analytical correlation with mass cytometry. We quantified iron and calcium mass/cell for murine T-lymphocytes with 3% and 8% 2-sigma intra-precision, respectively, when assessing thousands of cells/minute. T-lymphocytes exposed to a 625-fold difference in extracellular iron concentrations maintained close iron homeostatic control, varying ∼20% in iron content. Nevertheless, this relatively small variation strongly correlated with changes in cellular activation characteristics measured by flow cytometry. We also assessed human B-cell iron content, which was ∼10-fold higher than murine T-lymphocytes. Overall, we demonstrate rapid iron quantification at single-cell level in different cell types and relate cellular iron content to cell function.

**Teaser:** Precise and rapid iron metallomics of lymphocytes by single cell ICP-MS is a powerful approach for accessing signatures of immunological status.

## Introduction

Iron is crucial for multiple cellular biochemical activities(*1*). The failure to maintain iron homeostasis has important consequences for human health and vitality(*2*). For example, hereditary haemochromatosis results in iron accumulation, leading to tissue and organ damage(*3–5*). Iron deficiency anaemia (IDA) is recognised as the world’s most common micronutrient deficiency, affecting over a billion people worldwide (*6*). Furthermore, the severity and prevalence of IDA is particularly felt in many low-middle income countries, where a combination of malnutrition and endemic diseases give rise to unfavourable fates of disease pathogenesis, neurological developments and mortality(*7*).

IDA can also hinder the functioning of the immune system with consequences for health, including vaccine response(*8*). Stable homeostatic control of iron is crucial for effective functioning of T-cells and B-cells, where iron uptake is essential for cellular proliferation(*9*, *10*), activation(*11*) and metabolic reprogramming(*12*).

Iron deficiency is typically characterised by measurement of plasma parameters such as ferritin, or transferrin saturation, which act as proxy assessments of iron stores and iron availability for cells, respectively. However, the manifestations of iron deficiency depend on the iron content of cells and how cellular functions are altered by lack of iron. Our knowledge of how iron is distributed between different cell types and how cells respond to different levels of iron availability is currently limited(*13*). This largely reflects the inherent analytical difficulties in accurately measuring small variations in cellular iron levels amongst representative populations of cells.

Currently, cellular iron contents are typically determined by bulk dissolution, followed by analysis of the resultant solutes by a variety of atomic spectroscopy techniques (for example, inductively coupled plasma atomic emission spectrometry(*14*)). Although appropriate for determination of trace element contents, bulk approaches such as these are incapable of revealing any heterogeneity amongst cell populations. This, coupled with appreciable errors arising from cell counting methods, may lead to large uncertainties in the derived mass per cell values.

Analyses of individual cells by microanalytical methods, such as nanoSIMS and synchrotron µXRF, can enable the elemental quantification on a per-cell basis.

However, these microanalytical approaches are limited by the number of cells that may be analysed, the fixing procedures required and, in the case of synchrotron-based instruments, timely access to the instrument. Although they may deliver an accurate measure of cellular elemental content, the small number of cells that can be realistically analysed hinders estimates of intra-population variation.

An alternative approach to quantify the iron content of individual cells is to employ single cell inductively coupled plasma mass spectrometry (SC-ICP-MS). This technique relies on aspirating a solution containing a suspension of cells into an ICP- MS, such that they are dispersed sufficiently to be ionised, and subsequently analysed, sequentially. The ICP-MS instrument is operated in time-resolved mode, where analytical time intervals are set sufficiently short that cells transiting through the instrument can be discriminated from the background and ion counts arising are integrated to yield a total cellular mass for the element of interest. The SC-ICP-MS approach allows for the analysis of individual cells at a frequency of hundreds to thousands of cells per minute, sufficient for exploring cell population heterogeneity and providing a potentially rapid screening tool. However, measurement of iron by this analytical method has been successfully described in only a very few research studies. Upon examination of the literature and to our best knowledge, only one study has successfully quantified single cell iron by SC-ICP-MS (specifically in Raji cells for chemotherapy-drug uptake(*15*)), in addition to two others which have quantified iron levels in bacteria(*16*, *17*).

In this study, we optimised SC-ICP-MS for cellular material and cross-validated data against both ‘traditional’ bulk ICP-MS and CyTOF. We assessed single cell metallomics in populations of three different types of lymphocyte (Jurkat cells, OT-I murine T-cells and human primary B-cells). We measured iron in murine T-cells exposed to iron sufficient and iron depleted conditions and related iron content to cell activity.

## Results

### Single cell metallomics by SC-ICP-MS validated by titrated intercalation experiments

Rhodium and iridium, in their 3+ cationic forms are very effective metallointercalators when complexed with polycyclic aromatic ligands(*18*, *19*). They are often used in high-dimensional phenotypic/ functional single cell analyses by CyTOF mass cytometry for this reason (*20*). Moreover, as neither of these metals are normally utilised in biology, are both chemically rare (thus unlikely to reveal any contamination issues), and in their ionic form generate highly sensitive signals by ICP-MS, they can also be used to validate SC-ICP-MS as a metrological metallomic analytical technique.

In this experiment, titrated concentrations of rhodium and iridium intercalators were added to Jurkat cells, a prototypical T-cell line, and uptake was assessed in the attogram/cell (10^-18^g) mass range by SC-ICP-MS, Mass Cytomtery (CyTOF) and bulk ICP-MS (see Fig. 1A). PerkinElmer’s Syngistix Single Cell software module was used for realtime viewing of the single cell metallomics and post-measurement evaluation of SC-ICP-MS analysis, where the peak area from each cell event was used to determine the mass of metal per cell. The PerkinElmer NexION 350D ICP-MS was used to measure the rhodium-intercalated cells, and the PerkinElmer NexION 5000 for the iridium-intercalated cells.

**Fig. 1.**
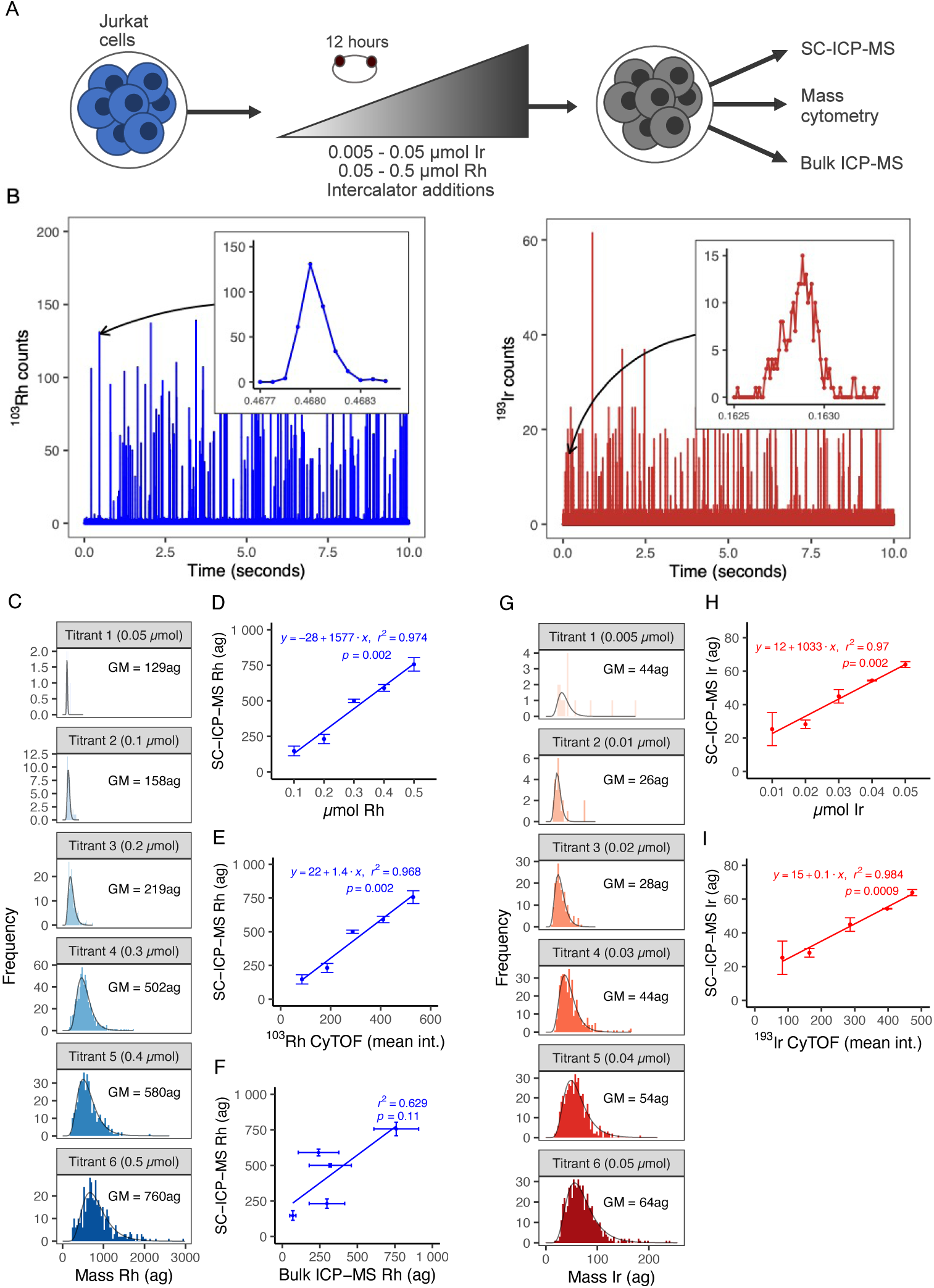
Validation of SC-ICP-MS from rhodium (blue) and iridium (red) intercalation experiments. **(A)**. Experimental design for validation of SC-ICP-MS by a metal intercalation approach; **(B).** Excerpts of time-resolved mass spectra of intercalated Jurkat cells, intercalated with 0.3µmol rhodium (left) and 0.04µmol iridium (right). Each spike represents a metallomic event, where their individual areas correlate to mass of rhodium or iridium within each measured cell. The metallomic plumes from each metal take ∼0.0004 – 0.0005s to transit through the instrument (shown in the inset peaks). The inset peaks also illustrate the resolutions achieved for the event profiles, where measurements were taken at intervals of 75µseconds (rhodium) and 10µseconds (iridium); **(C).** Mass-frequency histograms presenting single cell metallomic distributions from the rhodium intercalation series (GM = population geometric means). Mass quantified in attograms (ag) (10^-18^g); **(D-F).** Correlation plots of SC-ICP-MS (geometric means from rhodium experiment) versus intercalation molarity, mean CyTOF intensities and bulk ICP-MS analysis, respectively; **(G).** Mass-frequency histograms presenting single cell metallomic distributions from the iridium intercalation series (GM = population geometric means); **(H-I).** Correlation plots of SC-ICP-MS (geometric means from the iridium experiment) versus intercalation molarity and mean CyTOF intensities, respectively. Error bars shown in all correlation plots present 2-sigma precision for all SC-ICP-MS data points [n=4]. Error bars from bulk ICP-MS comprise a combination of ICP-MS and cell counting uncertainties, which are combined using the product rule.

The single cell metallomic approach using ICP-MS (as SC-ICP-MS) allows each cell from a population to be sequentially introduced to the instrument, where they are individually vapourised, atomised, and their metallomic constituents ionised for measurement within durations of around 200-400 µseconds(*21*). Known as a “cell event”, it is essential the instrument is able to capture the measurement of the metallomic plume for a particular element during its transit through the mass spectrometer. In this study we harness the high-frequency scanning capability of the PerkinElmer NexION ICP-MS to measure metals as mass-spectral fingerprints with minimal detector exposure times (dwell times), at intervals that are much lower than cell event durations (up to frequencies of 100,000Hz). This enables us to rapidly collect highly precise single cell metallomics, and to measure hundreds of cells per minute.

Fig. 1B present excerpts of realtime metallomic data taken from the rhodium and iridium SC-ICP-MS analyses of the intercalated cells, where each individual spike represents a single cell event. The peak insets demonstrate the high level of detail per cell event, where the beginning, end and apex of the example events are clearly defined (see insets within Fig. 1B). In the literature it has been postulated that low detector dwell times can adversely affect SC-ICP-MS data quality(*22*), primarily from compromised signal intensities. However, from our data we instead found that the detailed peak profiles, in addition to extremely low background baselines, conversely enhanced analytical figures of merit (transferable metrics for analytical performance (*23*)), such as linearity and precision. Moreover, we prove that high-frequency scanning also provides the prospect of filtering out fragmented cells versus real cell events, where the former transit through the instrument from distinguishably shorter event times than intact cells (see Fig. S1 in supplementary text).

Mass frequency histograms from the rhodium and iridium SC-ICP-MS measurements of each intercalated condition are shown in Figs. 1C and 1G, respectively, which are presented with optimised bin-sizes and log-normal fittings. Background corrections were also applied, which were defined by eliminating all cell events that fell short of the signal duration of a well-defined peak from the mass spectra (see supplementary materials for further information). Following such corrections, clear successions in metal mass per cell are found with increasing intercalation concentration, revealing the effectiveness of SC-ICP-MS to measure metallomic profiles of entire cell populations throughout 10^-18^g mass ranges. Heterogeneity of cellular metal uptake is also well-defined within each population, where interquartile ranges of 48ag (Titrant 2/ 0.1µmol) to 385ag (Titrant 6/ 0.5 µmol) were found from the Rh intercalation experiment; in addition to 7.5ag (Titrant 2/ 0.01 µmol) to 36ag (Titrant 6/ 0.05 µmol) from the Ir intercalation experiment (see Table S1 in supplementary text).

Additionally, the linearity in single cell mass progression throughout both titrations is clearly demonstrated from the clear linear regressions, significant r^2^ values (>0.97) and p-values (0.002) exhibited in Figs. 1D and 1H, where scatter plots of the population geometric means against the titrated intercalator molarity are presented.

Duplicate and triplicate aliquots from each condition of both titrations were measured by CyTOF and bulk ICP-MS to compare signal outputs. Excellent agreements were found between SC-ICP-MS and CyTOF from both titrations, which are presented in Figs. 1E and 1I as scatter plots of mean datapoints found from each sample (geometric means of metallomic distributions gained from the SC-ICP-MS and arithmetic means from CyTOF), where r^2^ >0.96 and p-values ≤0.002 are shown in both correlations. A similar correlation plot comparing SC-ICP-MS to bulk ICP-MS is also presented in Fig. 1F from the rhodium intercalation experiment, where on this occasion a much weaker correlation is displayed (r^2^ = 0.63, p-value = 0.11).

The contrast in analytical performance found between the single cell and bulk analytical techniques tested in this experiment is driven by the considerably enhanced detection capability from the former methods, where the combination of better fittings to the linear regression model (revealed from the enhanced r^2^ values) and the attainment of highly significantly p-values (<0.05) from each titration are found (see Figs. 1D, 1E, 1H and 1I). This is further emphasized by the bulk ICP-MS results gained from the lower-level iridium titration, where most conditions were found below the limit of quantification (thus not presented in Fig. 1).

### Calcium in T-cells is effective for determining transport efficiency for SC-ICP- MS

CD8+ T-cells were extracted from three OT-I mice. Fig. 2A presents an excerpt of the realtime single cell iron output from the T-cells analysed in this experiment, where akin to the previous rhodium and iridium intercalation studies, high frequency scanning was employed to capture high-definition profiles of each single cell ionic plume event.

**Fig. 2.**
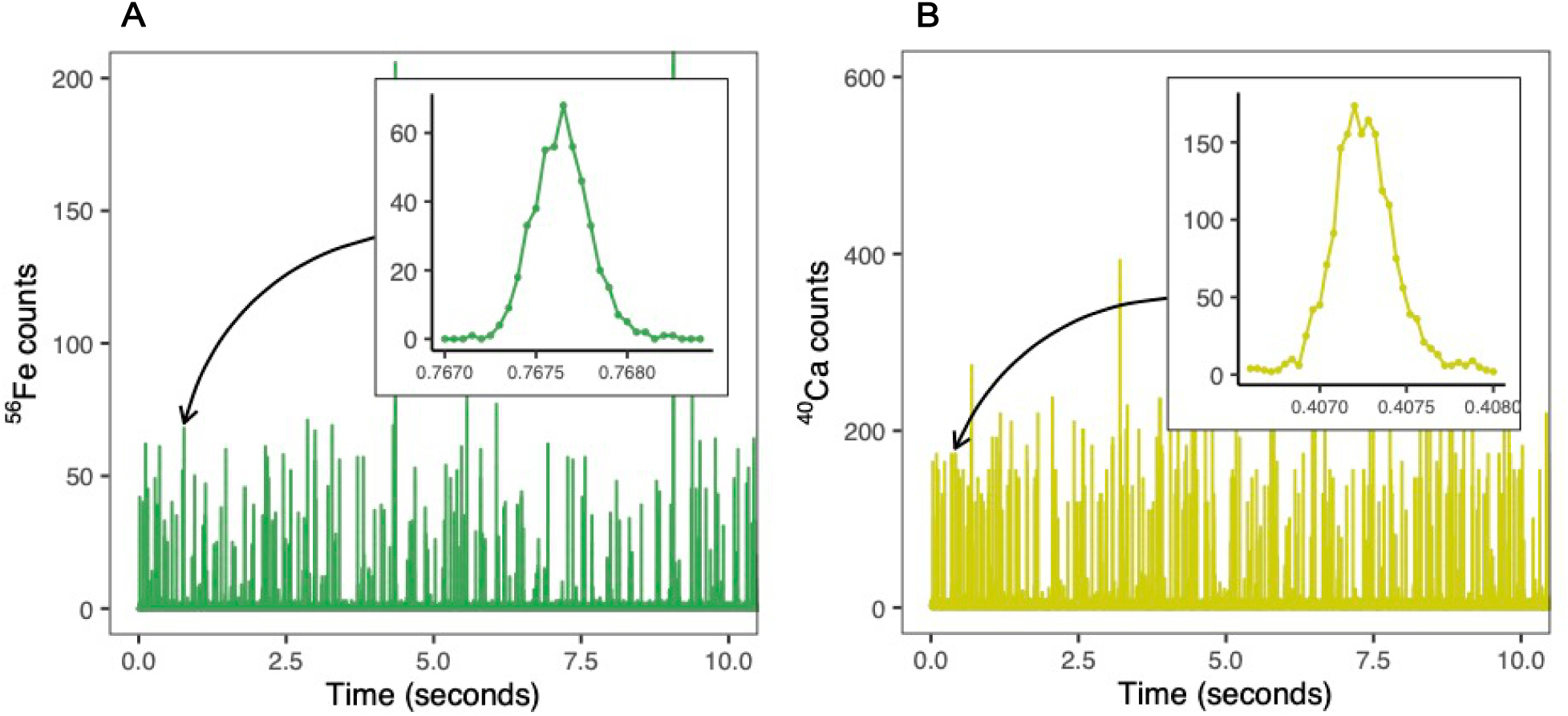
Time-resolved single cell mass spectra measured by SC-ICP-MS from murine T-cell experiments. **(A)**. An excerpt of time-resolved mass spectra from iron SC-ICP-MS analysis of T-cells taken from mouse 3 and the 0.005mg/mL condition; **(B).** An excerpt of time-resolved mass spectra from calcium SC-ICP-MS analysis of T-cells taken from mouse 3 and the 0.025mg/mL condition. Each spike in both **(A)** and **(B)** represents an individual metallomic event, where their individual areas correlate to mass of calcium or iron within each measured cell. The metallomic plumes from each metallomic event take ∼0.001s to transit through the instrument (shown in the inset peaks in both **(A)** and **(B)**). The inset peaks also illustrate the resolutions achieved for the event profiles, where measurements were taken at intervals of 40µseconds (calcium) and 50µseconds (iron). All measurements conducted using the PerkinElmer NexION 5000 ICP-MS.

Additionally, we also used SC-ICP-MS to rapidly scan for calcium within the same cell suspensions (see Fig. 2B for an excerpt of realtime calcium metallomic output). In lymphocytes, calcium is an endogenous element that is contained within a similar mass range to iron (circa. one magnitude higher)(*24*). Coupling the measurement of calcium metallomics to this experiment provided a direct means of determining the transport efficiencies of cell metallomic detection – the proportion of cells entering the plasma and being detected by the ICP-MS over the cell concentration in the measured aliquot, by the particle frequency method(*25*). Moreover, as the calcium mass spectra returned profiles that were constrained within tight mass ranges and similar duration times, it was a very useful proxy for determining a true value of cellular transmission and detection. From these data we determined a mean calcium-derived transport efficiency value of 12%, which is within a similar range to other studies who also reported using endogenous elements as a proxy for transport efficiency(*15*, *26*, *27*). Furthermore, this accurate representation of cell transmission is also complemented with high precision figures of merit, which was demonstrated by the return of excellent signal to background ratios. This was not only attained by scanning directly on calcium’s major isotope (^40^Ca) through the coupling of the instrument’s dynamic reaction cell (DRC) and tandem mass spectrometry capability, but also through the utilisation of high purity Maxpar^®^ Cell Acquisition Solution Plus for an extremely low background (see Table 1 and Methods for details). The inset peak within Fig. 2B illustrates this, where signal intensities captured from the apex of the peak from this particular cellular event, were over 30 times higher than the average baseline readings adjacent to the peak [n=17].

**Table 1.**
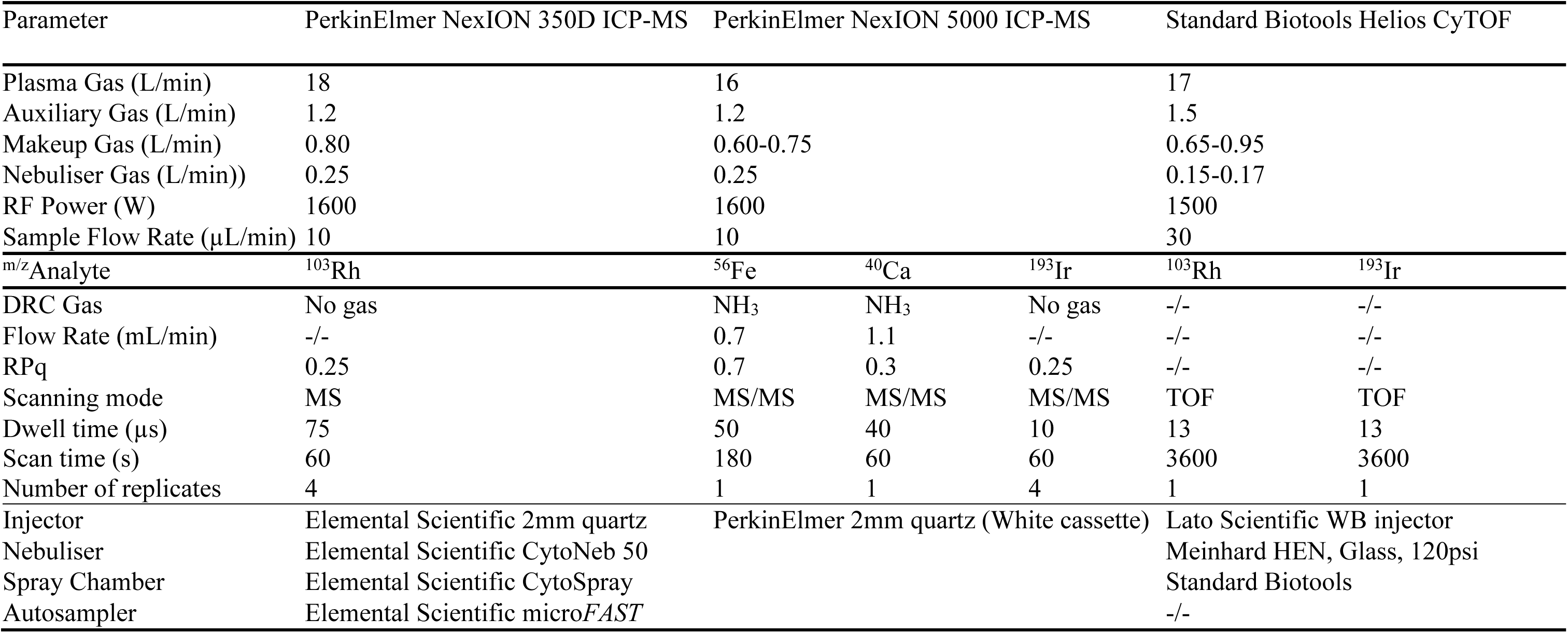
Operating conditions for SC-ICP-MS and CyTOF.

The single cell iron data also presents similarly tightly constrained cell event populations (for example as shown in Fig. 2A), but in contrast to the calcium metallomic data sporadic high-mass events were also present in the mass spectra profiles – which were most likely oxidised iron precipitates (as shown in Fig. 2A by the see peaks >200 counts in amplitude at ∼4 seconds and ∼9 seconds). However, from our high frequency scanning methodology, we were able to differentiate these discrete events from the data by their distinctly higher peak heights and widths.

Additionally, as they also lacked any defined gaussian/ lognormal distributions and consisted of values away from the well-defined cell distributions, threshold corrections from the cell distributions was relatively simple. After subsequent comparative filtering of the iron quantification data for only cell events, the post-filtered iron metallomic data presents a mean transport efficiency that is only 1% higher to the value obtained from calcium, at 13%, thus proving a highly accurate filtering method. Table S2 in supplementary text provides data of the transport efficiency derived for calcium and iron (after filtering) for each sample, in addition to similarly calculated transport efficiencies derived from the measurement of holmium scanning from EQ^TM^ Four Element beads. In contrast to cells, these polystyrene beads were resistant to lysing during the introduction phase of measurement, which is highlighted by the higher values, where an average transport efficiency of 27% was attained (see Table S2).

### Iron analysis by SC-ICP-MS requires chemical resolution by MS/MS for accurate detection

Since its inception in the 1990s, the DRC within PerkinElmer ICP-MS instruments (in addition to other similar manufactured reaction cells), has revolutionised the ability to accurately measure those elements that are affected by polyatomic interference (*28*, *29*). Its ability to negate polyatomic interferences, either by adopting exothermic reactions from reactive gases to either neutralise/ disassemble such ions, or by kinetic energy modulation/ dissociation from highly pressurised inert gases, has elevated analytical performance for a plethora of elements, including iron. However, such gases also sustain impedances (although tempered) to the analyte ions, which can complicate SC-ICP-MS analysis. This effect causes time-elongation of ion plume events and thus peak tailing, up to 6ms in the time-resolved spectra(*30*), which can detrimentally impact upon the accuracy of the resultant metallomic/ nanoparticle findings(*15*, *31*). Additionally, it can also increase the probability of recording doublets, especially when using high flow rates of NH_3_ as the cell gas for mass-shift scanning methods(*15*). We mitigated this impact in our experiments, by using a moderated flow of NH_3_ in the reaction cell, elicited by an MS/MS on-mass scanning approach for analysis – a chemical resolution method that still remains effective at removing such problematic spectral interferences, but one that does not inflict such significant impacts on the ionic energies(*15*, *30*). Here we observed only minimal elongations to such cell event data, where ∼1ms peak widths were reported by chemical resolution (iron), versus circa. 0.5ms when no cell gas was used (rhodium) (see Figs. 2A and 1B, respectively). This magnitude of protraction is 6-times lower than those reported from the mass-shift approach(*30*), engendering a negligible probability of recording doublets (i.e. from the occurrence of two simultaneous peak events).

To quantify uncertainty from our measurement method we examined analytical figures of merit, including precision and accuracy. For precision, determinations from consecutive measurement repeats from one of the samples measured [n=4] were evaluated, which revealed 2-sigma variability (as 2 times relative standard deviation) of 2.7% for iron and 8.3% for calcium. Additionally, repeat measurements of EQ^TM^ Four Element Calibration Beads for holmium (^165^Ho) at intervals throughout the analytical run [n=4] presented similarly-calculated precision values of 5.9%. As there are no certified reference materials available for this field of analysis, analytical accuracy was instead determined by measurement of an iron quality-control standard solution. This presented relative errors of 3.1% for iron and 8.1% for calcium.

### Cellular iron uptake in OT-I T-cells is limited by proliferation

OT-I T-cells derived from three individual mice were activated for 48 hours in iron-free media supplemented with titrated concentrations of holotransferrin (transferrin protein with two iron atoms bound) from 0.001mg/mL to 0.625 mg/mL, which characterised a range of conditions from severe iron deficiency to iron replete (see Fig. 3A). Mass frequency histograms of the background corrected single cell iron data for the titrants from each mouse are presented in Fig. 3B, which are presented with optimised bin sizes, log-normal fittings and annotated geometric mean values for each population. We observed only small increments in the geometric means of cellular iron content in populations exposed to increasing amounts of holotransferrin. These extrapolated average iron contents per cell (for each condition and each mouse), were subsequently plotted against each media iron concentration, where clear correlations are presented (see Fig. 3E). Although the geometric means of atomic iron content per cell only increased slightly (∼20%) over a 625-fold difference in holotransferrin, this indicates a general maintenance of cellular iron homeostasis in the face of a range of extracellular iron availability.

**Fig. 3.**
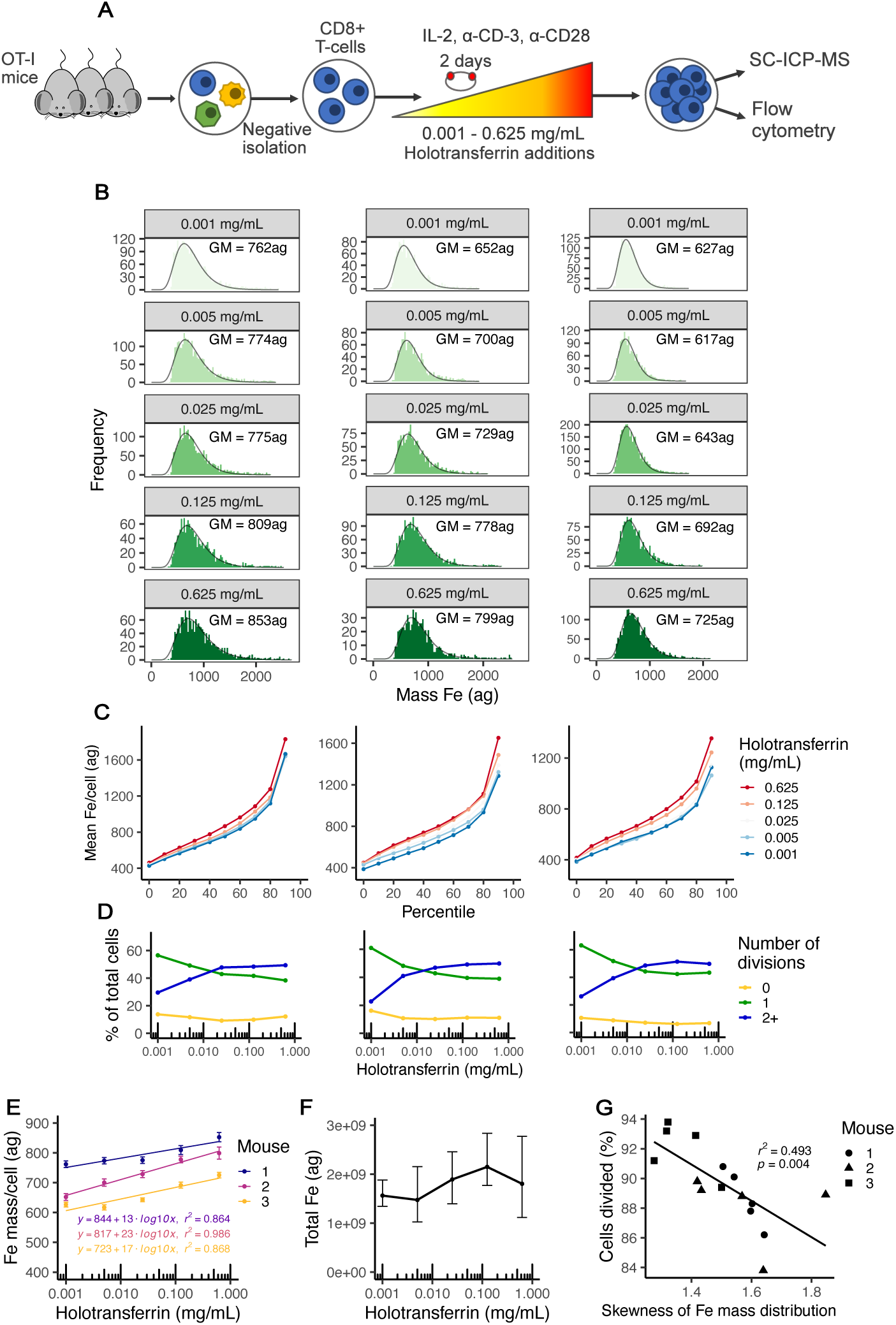
Results from murine T-cell iron deprivation experiment. **(A)**. Experimental design for the assessment of iron in murine T-cells that were exposed to iron media conditions ranging from 0.001mg/mL – 0.625mg/mL holotransferrin; **(B).** Histograms showing the mass-frequency distributions of iron mass/cell for each holotransferrin condition from cells taken from each of three mice. Mass distributions are shown in attograms (ag) (10^-18^g); **(C).** Line plots showing the mean iron mass/cell per tenth percentile for each holotransferrin condition from each mouse; **(D).** Line plots presenting proportions of cell proliferation activity per iron condition (percentage of total cells dividing 0, 1 or 2+ times) with log_10_ x-axes from each mouse; **(E).** Correlation plot presenting the geometric means from each distribution versus the iron condition for each mouse (with a log_10_ x-axis). Logarithmic trend lines, associated equations and r^2^ values are displayed for each profile. Error bars show the 95^th^ confidence intervals; **(F).** Line plot presenting the mean total amount of iron consumed by the cells from the three mice for each iron condition (with a log_10_ x-axis). Error bars show the range of total iron calculated per mouse. **(G).** Scatter plot presenting the inverse correlation between the magnitude of cell proliferation during the live culturing period against the skewness of the single cell metallomic distributions for iron.

Heterogeneity within the main distributions of each population was evaluated from their calculated interquartile ranges, where similar limited increases between the lowest and highest iron conditions are presented (see Fig. S2 in supplementary text). Equally, an evaluation of intra-population variability, in addition to diversity in iron-uptake behaviour at the tails of the populations was also determined upon segregating each population into percentiles, where means of each 10^th^ percentile were calculated and plotted per mouse in Fig. 3C. Between the 10^th^ and 70^th^ percentiles smooth trends are illustrated, reflecting well-constrained and tightly-defined log normal distributions in iron per cell (see Fig. 3B), which also transition in magnitude in accordance to their iron condition. However, beyond the 70^th^ percentiles, significant escalations in the average cellular iron levels are presented in every condition, which is replicated from cells isolated from each mouse. This observation, although characteristic of positively skewed distributions, indicates that the top 20% of cells within each condition contain discrete elevations in iron levels over the rest of the population. Furthermore, unlike the uniform transitions noted from the earlier percentiles, the rapid increases in average iron/cell within this range do not occur linearly, where distinctly higher elevations were found for the higher conditions (particularly 0.625mg/mL).

Unlike the previous intercalation studies, during the 2-day culturing period for this murine T-cell experiment, live cells were able to continually acquire available iron and proliferate (see Table S3 in supplementary text). To examine this effect, we analysed parallel aliquots of cell cultures by flow cytometry to assess the extent of cell division per condition for each mouse. The proportions of cells within each population that didn’t divide, divided once, or divided two or more times over the 48- hour period, are presented in Fig. 3D. The results present clear evidence of the escalation in proliferative activity in the higher iron-bearing conditions, particularly the proportion of cells undergoing 2+ divisions.

As mentioned above, although the geometric mean of iron content per cell from this experiment only varied by ∼20%, the total amount of cellular iron incorporated into the cell population is higher. Using live cell count data collected during cell harvest together with the geometric means of iron content/cell, it was possible to quantify the total amount of iron used by the cell populations in this experiment (see Fig. 3F). Of significance, the peak iron yield did not associate with the highest condition, which was instead found at the penultimate condition level (0.125mg/mL). This is likely a consequence of cytotoxic effects from the superfluous levels contained within the 0.625mg/mL condition, which is reflected by the lower live-cell counts obtained.

Furthermore, unlike the low levels of variability found in single cell iron levels between each iron condition, much wider differences were found in the total amounts of iron consumed per cell population, where a maximum variability of 94% was found between the highest and lowest calculated values.

The extent of proliferative activity was also found to correlate with the shape and tailing of each single cell iron population obtained by SC-ICP-MS, which was measured by their skewness values (see Fig. 3G). The degree of tailing, or skewness, in such distributions is also proportional to the amount of heterogeneity in the cell populations, thus revealing crucial detail on the spread of metallomic masses, or even potentially phenotypical variability. Across the culture conditions from this experiment, the skewness of the single cell iron mass distributions inversely correlated with their degree of proliferation (proportion of cells divided), where those populations comprising elevated skewness also posed enhanced phenotypical variability – affirmed by the associated increase in non-dividing cells that must accumulate higher levels of iron within the population (see Fig. 3G). Furthermore, all of the datapoints from mouse 3, with the exception of one, contain the lowest skewness values, together with the highest levels of cell division, which could explain why lower geometric means of the single cell iron distributions were presented from this mouse (see Figs. 3E and 3G). Also, as the cell division findings provide a high degree of correlation to the single cell iron data from the ICP-MS, we can discount any possibility of the high iron mass per cell datapoints being recorded as doublet cell measurements.

### Single cell iron metallomic data correlates with surface glycoprotein markers

In addition to measuring cell proliferation activity, flow cytometry was also employed to assess cell surface expression of CD71 (transferrin receptor) and CD25 (IL-2 receptor) per condition for each mouse. The data collated from these measurements are compared to the geometric means of iron mass from each cell distribution measured by SC-ICP-MS, where such data is presented as scatter plots in Figs. 4A and 4B, respectively. CD71 provides the primary uptake mechanism for transferrin-bound iron into eukaryotic cells. Notably, mutations that disable CD71-mediated iron acquisition cause immunodeficiency and impair proliferation of T-cells(*32*). CD71 is normally highly expressed by activated T-cells, but its synthesis is also regulated by intracellular iron content, with relatively iron deficient cells expressing higher levels of CD71 in order to more efficiently capture any available extracellular iron(*12*). We observed a clear inverse correlation between geometric mean iron content per cell and mean fluorescence intensity of CD71 (Fig 4A), providing an orthogonal assessment of cellular iron content downstream of cell-intrinsic iron sensing mechanisms.

**Fig. 4.**
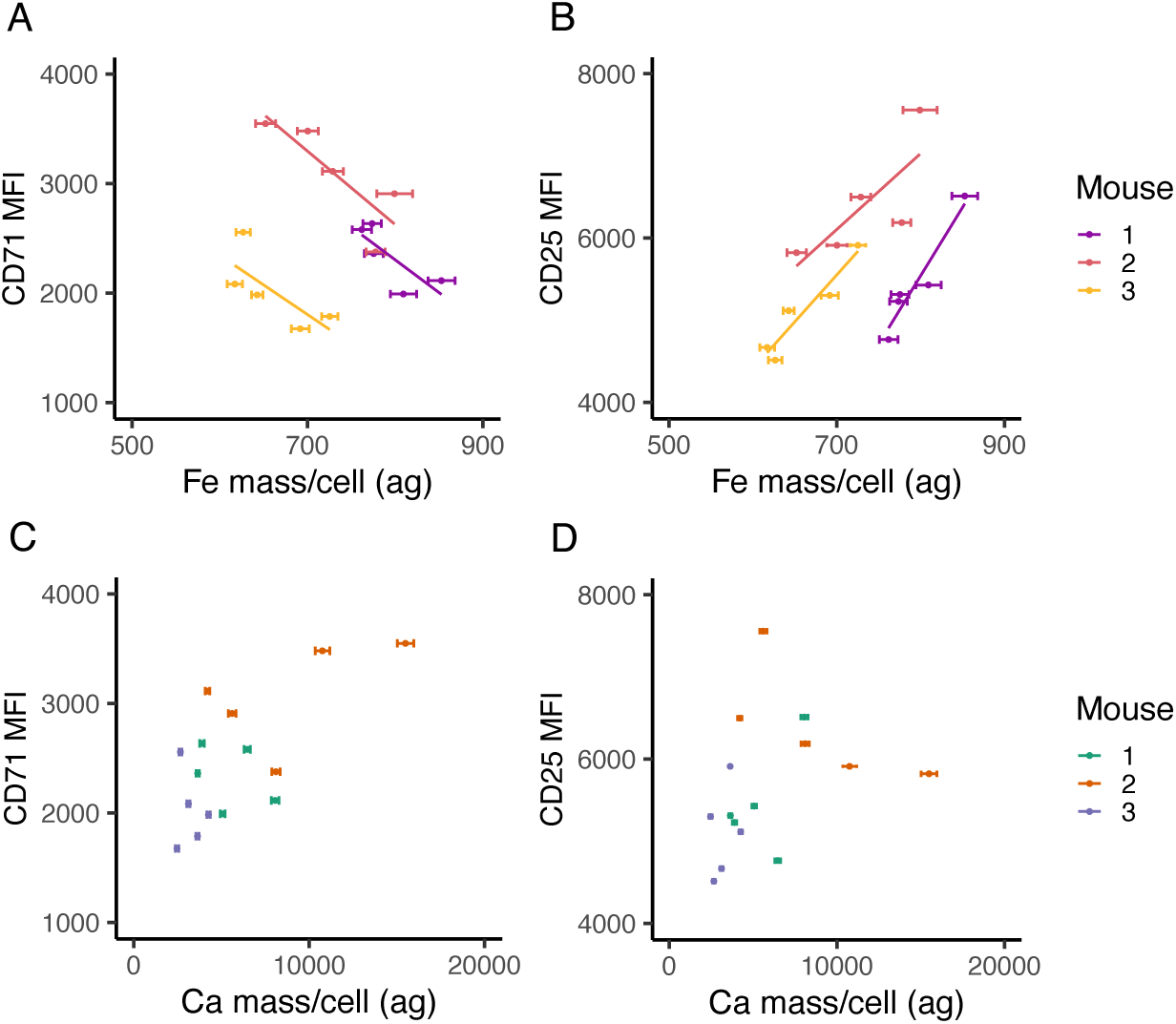
Correlation of surface protein marker expression determined by Flow Cytometry against SC-ICP-MS derived single cell metallomic contents. **(A)**. Mean fluorescence intensity (MFI) of CD71 (transferrin receptor) versus the geometric mean single cell iron content (ag) for each population distribution, from each holotransferrin condition; **(B).** Mean fluorescence intensity (MFI) of CD25 (alpha chain component of the IL-2 surface receptor on T-cells) versus the geometric mean single cell iron content (ag) for each population distribution, from each holotransferrin condition; **(C).** Mean fluorescence intensity (MFI) of CD71 versus the geometric mean single cell calcium content (ag) for each population distribution, from each holotransferrin condition; **(D).** Mean fluorescence intensity (MFI) of CD25 versus the geometric mean single cell calcium content (ag) for each population distribution, from each holotransferrin condition. Error bars present the 95^th^ confidence intervals for each metallomic distribution determined by SC-ICP-MS.

CD25 is the alpha chain component of the IL-2 surface receptor on T-cells. IL-2 signalling via CD25 promotes T cell growth and facilitates their differentiation after activation(*33*). We found that CD25 expression positively correlated with geometric mean iron content, in line with the importance of iron acquisition for cellular activation and growth, as also observed by the correlation of increased cellular iron and cell proliferation observed in Fig. 4B.

As a comparison, we plotted expression of surface protein markers to the geometric means of calcium mass from each cell distribution measured by SC-ICP-MS. Figs. 4C and 4D present the associated scatter plots, with no correlations of calcium content with either CD71 or CD25, showing the relative specificity of the relationships between iron content and T-cell activation.

### Precise single cell iron mass distributions from human primary B-cells

To move beyond murine systems and analyse human cells, we examined primary B-cells extracted from the peripheral blood of three healthy donors. After 3 days of in-vitro culturing in R10 media the cells were measured by SC-ICP-MS, using a similar method used for the measurement of the previous murine T-cells. B cell purity was assessed through analysis of CD19 surface protein expression.

An excerpt of the realtime iron metallomic output are presented in Fig. 5A, where unlike the murine T-cell experiment, peaks comprising wider size ranges were obtained. This observed increase in heterogeneity likely indicates increased phenotypical variability in cellular iron from this cell type. In a similar fashion to the murine T cell analyses, background corrections were applied to remove the presence of ionic and fragmented cell artifacts, with transport efficiencies ranging from 30% to 52%.

**Fig. 5.**
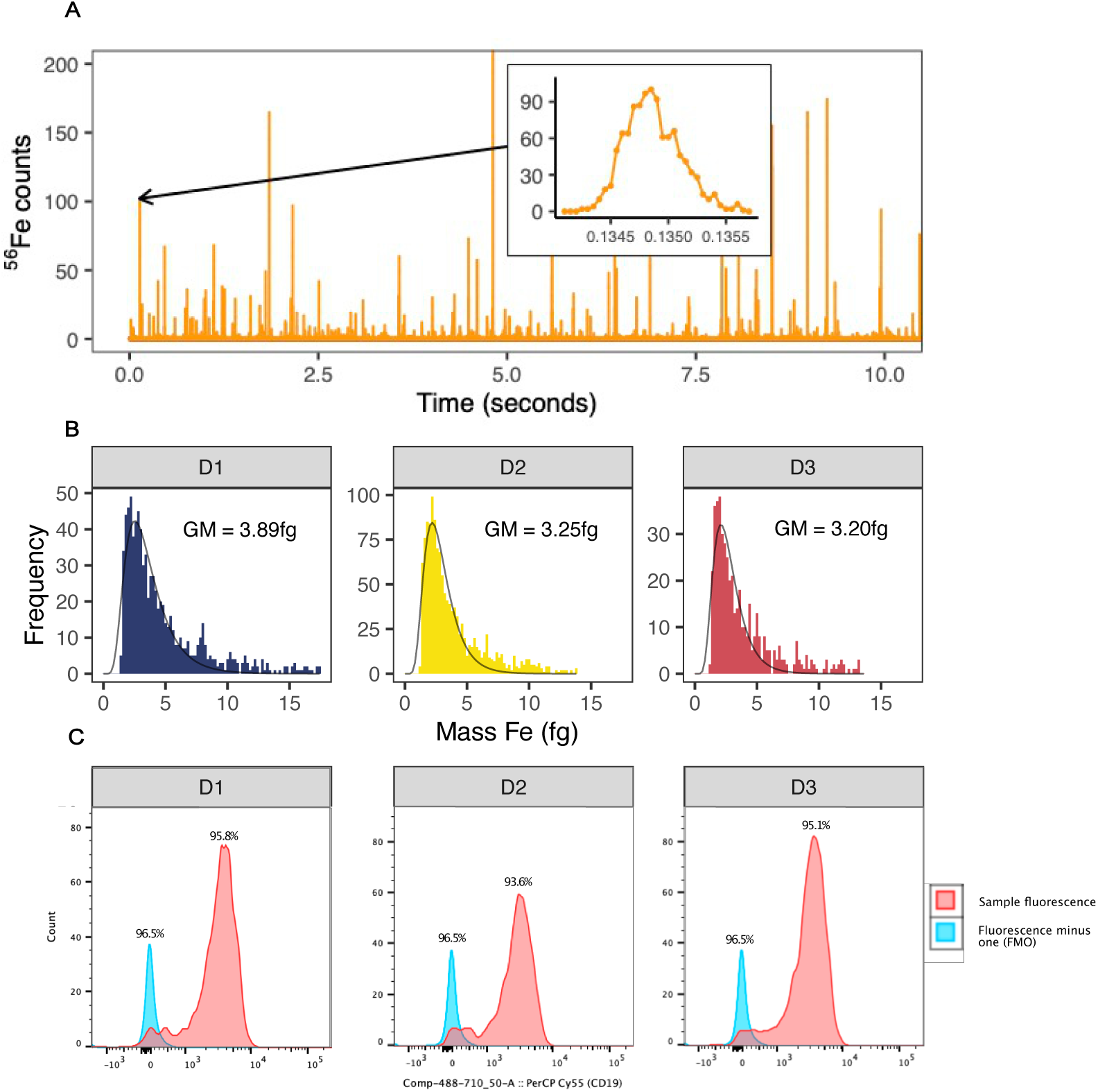
Results from primary human B-cell experiment. **(A)**. An excerpt of time-resolved mass spectra measured for iron by PerkinElmer NexION 5000 SC-ICP-MS from human B-cells taken from donor D1. Each spike represents a cell metallomic event, where their individual areas correlate to mass of iron within each cell. The metallomic plumes take ∼0.001s to transit through the instrument (shown in the inset peak). The inset peaks also illustrate the resolutions achieved for the event profiles, where measurements were taken at intervals of 50µseconds. **(B).** Mass frequency histograms presenting distributions of single cell iron content determined by SC-ICP-MS for each of three donors, D1-D3, respectively. Mass distributions are shown in femtograms (fg) (10^-15^g). GM = distribution geometric mean for each population. **(C).** Histograms of CD19+ cell counts, determined by flow cytometry, for each of three donors, D1-D3, respectively.

Mass frequency histograms of such corrected data for each donor (D1 – D3) are shown in Fig. 5B, which are presented with optimised bin sizes, log-normal fittings and annotated geometric means for each population. Similar mass ranges were found for each sample, where 2-sigma variability was 0.77fg between the three samples.

Additionally, wide distributions are presented, which reflects the wider range in peak sizes noted in Fig. 5A, but extrapolation of the mean values, together with the above uncertainty value presents overall metallomic values that are within range to similarly measured Raji B-cells(*15*). Moreover, the complimentary CD19 provided confirmation of B cell purity in our measured cell aliquots, where recoveries ranging from 93.6% - 95.8% were reported (see Fig. 5C).

## Discussion

One of the main advantages of SC-ICP-MS for metallomic analyses is its detection capability, where it is able to quantify metals encapsulated within individual cells throughout attogram mass range (10^-18^g)(*34*). Additionally, unlike other solution-based analyses, dilutions applied to reduce the cell concentration never impact on the signal intensity of the cell events, where instead it may help to reduce the dissolved baseline and thus improve signal: noise ratios. In further attempting to achieve higher counting statistics and reduced data volumes, there is a temptation in many SC-ICP-MS studies to collate metallomics from dwell times that exceed the transit time of a cell event (circa. 200-400 µseconds(*21*)). However, we present crucial findings that instead advocate higher scanning frequencies, particularly at intervals much faster than cell events. Nevertheless, it is important to mention that the ability to scan at frequencies greater than the transit times of cell events may be a limiting factor of the particular ICP-MS instrument being used (*35*, *36*). Several studies have interrogated SC-ICP-MS for the highest achieving dwell times for metallomic analyses(e.g. (*22*, *35*)), with the outcome often suggesting the adoption of the shortest available interval time. In addition to negating the possibility of recording doublets, reduced backgrounds were particularly favoured from such measurement intervals. Moreover, in our study we also emphasize the necessity to scan cell events at high frequency intervals to also provide elucidation between “real” cell events and cell fragments/ debris, which in this study, particularly enabled the accurate detection of iron in our murine T-cell experiments.

SC-ICP-MS does have its limitations though, notably regarding transport efficiency, where components of the sample introduction system can cause significant cell losses. Nevertheless, through the coupling of a dedicated single cell microflow injection system, we were able to constrain such lysing effects and consistently achieve mean cell-derived transport efficiencies of nearly 15% (see Table S2 in supplementary text). Additionally, the proficient nature of SC-ICP-MS to examine heterogeneity within cell populations may not be a requirement for all fields of metallomic research, where bulk approaches providing mean values may otherwise suffice (e.g. (*37*, *38*)).

Nevertheless, as we discovered from the method evaluations, inaccuracies from cell counting can inhibit overall accuracy when adopting this approach.

In this study we provide a fully-validated evaluation of our SC-ICP-MS method from the intercalation experiments, followed by applied murine T-cell and primary B-cell experiments. Through the careful formulation of titrated rhodium and iridium metal- intercalated cells, we present the clear capability of our analytical methodology to precisely quantify cellular metal uptake throughout attogram/cell mass range (as described above and shown in Fig. 1). Heterogeneity throughout the main distributions was also evidently captured and evaluated from their interquartile ranges, where fluent progression from the lower (0.005 µmol – 0.05 µmol) Ir, to the higher (0.05 µmol – 0.5 µmol) Rh intercalation experiments was attained (see Table S1 in supplementary text).

From in-depth evaluations of the murine T-cell experiment, not only did we observe equivalence in heterogeneity throughout the main population distributions to the intercalation experiments, but also revealed evidence of phenotypical variability within the uppermost percentiles of each population. Significantly, the magnitude of iron mass per cell within this sub-population also escalated with iron condition, suggesting disparities in metal uptake behaviours associated with iron status.

Furthermore, we also correlated the skewness of each distribution to the magnitude of proliferation, where a distinct inverse correlation presented direct evidence of the influence that the magnitude of proliferation had on the overall heterogeneity of the population. The geometric means of each population provided comprehensive broader-scale assessments, where moderated uptake profiles were found for each mouse - ascertained from only ∼20% increases observed. This occurred even though the media concentrations were titrated throughout a range that varied by a factor of 625 times. From this we elucidate the overall influence that metabolic activity played during utilisation of bio-available iron from the live culturing, providing the limited iron per cell uptake profiles that we found. Moreover, this relationship replicates findings from a similar iron culturing study, where magnetotactic bacteria, rather than lymphocytes, were examined for their metabolic response to varying extracellular iron conditions (*16*). The logarithmic iron uptake trends presented in (*16*) are analogous to those presented here, which provides encouraging confirmation of the results that we have found. Finally, we provide validation data towards our SC-ICP-MS metallomic findings from glycoprotein markers measured by flow cytometry (CD71 and CD25). Both sets of results provide significant correlations that provide significant direct evidence of metabolic activity to downstream iron contents.

Overall, these data findings show the powerful application that SC-ICP-MS can offer in metallomic research, where its ability to rapidly and precisely scan the profiles of entire cell populations can reveal important physiological and/or biochemical behaviours in biological research. Nevertheless, as this is still an emerging field of research there are limitations to the results gained, specifically in the scope of background data filtering and the determination of ‘real’ cell results against fragmented entities. As described above we utilise an accurate and robust filtering approach by using the rapid scanning capability of the NexION ICP-MS in addition to ancillary calcium metallomic data to decipher between such data findings, but mathematical models capable of furthering such data corrections are required to reinforce such corrections.

## Materials and Methods

### Jurkat cell line, rhodium and iridium intercalation

Clone E6-1 Jurkats were purchased from ATCC; a clone of the Jurkat-FHCRC cell line (derivative of the original Jurkat cell line). They were cultured in R10 media (RPMI 1640 supplemented with 10% foetal bovine serum, 1% penicillin-streptomycin and 1% glutamine) and incubated at 37°C and 5% CO_2_ in T75 flasks. For the metal intercalation experiments, freshly passaged cells were divided into subsets (each containing cell concentrations of circa. 4x10^6^ cells/mL) and rinsed by centrifugation with Standard Biotools Maxpar^®^ phosphate buffered saline (PBS), prior to subsequent fixation in 4% paraformaldehyde for 10 minutes at room temperature. Titrated concentrations of either rhodium or iridium intercalators (Standard Biotools’ 500µM rhodium Cell- ID^TM^ or 125µM iridium Cell-ID^TM^, respectively) were then doped into each sample to form intercalation concentration ranges of 0.05 µM – 0.5 µM and 0.005µM – 0.05 µM, respectively. The cell samples were then stored overnight at 4°C to ensure complete penetration of the Cell-ID^TM^ organo-metallic compounds by passive diffusion through the permeated membranes of each cell. The following day each sample was divided into two to provide aliquots for both SC-ICP-MS and CyTOF analysis. Prior to analysis the cells were rinsed with Standard Biotools Maxpar^®^ cell staining buffer by centrifugation to remove any excess metal accumulation from the cell surfaces.

### Mice and T cell isolation from peripheral blood, iron titration and in-vitro proliferation

OT-I mice (2 x 12-week-old males, and 1 x 13-week-old male), were originally obtained from Audrey Gerard, University of Oxford, and were housed in individually ventilated cages. All animal work was completed under the authority of UK home office project and personal licenses under the Animals (Scientific Procedures) Act (ASPA) 1986. Mice were sacrificed via rising concentration of CO_2_ followed by cervical dislocation. Plates for CD8+ T-cell culture were pre-treated with 5 ug/mL α-CD3 (Biolegend, 100239) in phosphate buffered saline for 2 hours at 37°C. Spleen and lymph nodes were collected from euthanised mice and macerated through 40 μm filters using PBS supplemented with 2% fetal bovine serum and 1 mM EDTA (Invitrogen, AM9260G). CD8+ T-cells were isolated from the single cell suspension using the EasySep Mouse CD8+ T-cell isolation kit (Stem Cell Technologies, 19853) and the EasyEights EasySep magnet (Stem Cell Technologies, 18103). Isolated cells were stained with cell trace violet (CTV, Invitrogen, C34557) for 8 minutes at 37°C in PBS and then washed. CD8+ T-cells were plated at a concentration of 0.5x10^6^ cells/mL on the α-CD3 pre-treated plates. Cells were grown in iron free media (RPMI1640 (Gibco, 21875034), 10% iron free serum substitute (Pan Biotech, P04-95080), 1% glutamine (Sigma Aldrich, G7513-100ML) and 1% penicillin/streptomycin (Sigma Aldrich, P0781-100ML)) supplemented with set concentrations of holo and apotransferrin. Human holotransferrin (R&D systems, 2914-HT-001G) was added at concentrations of 0.001 mg/mL to 0.625 mg/mL. Total transferrin levels were kept at a constant concentration of 1.2 mg/mL by adding the appropriate amount of human apotransferrin (R&D systems, 3188-AT-001G). Cells were also treated with 50 μM β-mercaptoethanol (BME, Gibco, 31350-010), 1 μg/mL α-CD28 (Biolegend, 102115) and 50 U/mL IL-2 (Biolegend, 575402) to activate the cells. CD8+ T-cells were cultured at 37°C, 5% CO_2_ for 48h.

After incubation the cells were harvested, counted and aliquots divided between SC-ICP-MS and Flow Cytometry; where ∼ 2x10^6^ cells/mL were retained for SC-ICP-MS. The SC-ICP-MS cell aliquots were then rinsed twice by centrifugation with Standard Biotools Maxpar^®^ PBS, followed by resuspension in 1mL 4% paraformaldehyde for fixation at room temperature for 10 minutes. The samples were then rinsed twice by centrifugation with Standard Biotools Maxpar^®^ Cell Staining Buffer to remove any excess iron remaining from the cell surfaces, followed by resuspension in Standard Biotools Maxpar^®^ Fix and Perm reagent for overnight storage at 4°C.

### Human blood donations: B cell isolation, purification and in-vitro proliferation

Blood samples taken from three healthy donors from the John Radcliffe Hospital, Oxford, United Kingdom, were utilised for single cell iron analysis in B-cells by SC-ICP-MS. Each sample was collected after obtaining written consent and ethical approval from the University of Oxford’s Central University Research Ethics Committee (CUREC). The samples were collected in EDTA, which was followed by PBMC isolation by density gradient centrifugation: Greiner Bio-One Leucosep tubes, containing 15mL of Lymphoprep (Stem Cell Technologies) and collected blood were centrifuged at 1000 x g for 1 minute at ambient temperature. EDTA blood was extracted into the upper chamber of the Leucosep tube and centrifuged at 1000 x g for 15 minutes with no brake. The cloudy buffy-coat layer, containing PBMCs was extracted and the cells were subsequently rinsed twice with R0 media (RPMI 1640 supplemented with 1% penicillin-streptomycin and 1% glutamine) and PBS, respectively. CellTrace Violet (Thermo Fisher Scientific) was added to the rinsed PBMCs as a tracer for proliferation and incubated at 37°C in 5% CO_2_ for 8 minutes. After incubation the cells were rinsed with R10 media, counted and then diluted to 8x10^6^ cells per mL in R10 media. Two million cells were subsequently added per well into a 24-well rounded bottom plate, together with aliquots of 0.25mL of R10 media and 0.5mL of R10 media supplemented with 1µg/mL R848 (Stem Cell Technologies) and 10 ng/mL of recombinant IL-2 (PeproTech). The prepared cells were then cultured for 3 days at 37°C in 5% CO_2_. Following polyclonal stimulation, the cells were harvested, washed in R10 media and counted. B-cells were then purified from the other harvested PBMCs by negative selection using a Human B-cell Isolation Kit (Stem Cell Technologies) according to manufacturer’s instructions. Following purification, the B-cells were transferred to a 96-well rounded bottom plate and rinsed with PBS, following Fc Receptor (FCR) blocking and live/dead staining. This was followed by the subsequent labelling of the cells with combinations of anti-CD19- PerCP Cy5.5, anti-CD21 (Alexa Fluor 700), anti-CD27 (PE-Cy7), anti-CD38 (BV510), anti-CD69 (BV605), and anti-CD71 (PE/Dazzle 594) in PBS in addition to incubation for 20 minutes on water ice and fixation buffer (Biolegend). Prior to intracellular staining with anti-IgG (BV711) and anti-IgD (FITC) the cells were permeabilised with perm buffer for 20 minutes on water ice. Prior to measurement by Flow Cytometry, fluorescence minus one controls (FMOs) were included for each marker, in addition to an unstained control

### Single cell Inductively coupled plasma mass spectrometry (SC-ICP-MS)

Following all of the methodologies described above, the prepared cell suspensions were also rinsed a further three times in Standard Biotools’ Maxpar^®^ Cell Acquisition Solution Plus (CAS+) prior to SC-ICP-MS analysis. This was undertaken to ensure both the removal of any remaining residually-retained metals from the cell surfaces, in addition to an exchange into a suspension media suitable for analysis by this technique. Indeed, this reagent is proven to be an optimal choice for analysis utilising our analytical setup over other commonly used carrier reagents(*39*, *40*), where its combination of a neutral pH in addition to a higher ionic content than water provides a higher analytical performance for metallomic analysis when combined with a wider bore injector. Succeeding this rinsing protocol, the final suspensions were filtered through 35µm nylon mesh filters, and the subsequent cell suspensions counted, diluted to 10^5^ cells/mL cell concentrations (if required) and then immediately measured by SC-ICP-MS. Supernatants from the final rinse cycles of selected samples were also retained for measurement, to test for any leakage of intracellular metals.

All SC-ICP-MS measurements were conducted using either a NexION 5000 multi- quadrupole ICP-MS (PerkinElmer) or a NexION 350D ICP-MS (PerkinElmer), in time-resolved mode (see instrument conditions stated in Table 1). Both instruments were equipped with an Elemental Scientific Inc. single cell introduction system, which comprised of a CytoNeb 50 nebuliser, a CytoSpray linear pass spray chamber, a 2.0mm tapered injector (PerkinElmer White Cassette torch with 2.0mm injector for the NexION 5000) and a micro*FAST* autosampler (which provided an additional final agitation of the suspension using its ‘Mix’ submethod to ensure homogenisation of the aliquoted cell suspension for analysis). This apparatus, like many others also used for single cell metallomic research(*15*, *22*, *41*), was essential for this analysis to ensure the highest levels of cell transmission to the instrument, where micro-flow volume injections of cells were analysed for precise single cell metallomics by the mass spectrometer from discrete measurements of their resultant ionic plumes. Details relating to the optimisation of this instrument setup is described in supplementary information.

### Bulk Inductively coupled plasma mass spectrometry (bulk-ICP-MS)

Bulk ICP- MS analysis utilised cell aliquots remaining after the metal intercalation SC-ICP-MS analyses, where approximately 0.2x10^6^ cells were dissolved in 2% v/v HNO_3_ within metal-free centrifuge tubes at ambient temperature for 48 hours. Measurements were conducted using the NexION 350D ICP-MS (PerkinElmer) using instrument conditions described in Table 2.

**Table 2.** Operating conditions for bulk-ICP-MS.

| Parameter | PerkinElmer NexION 350D ICP-MS |
| --- | --- |
| Plasma Gas (L/min) | 18 |
| Makeup Gas (L/min) | -/- |
| Auxiliary Gas (L/min) | 1.2 |
| Nebuliser Gas (L/min)) | 0.915 |
| RF Power (W) | 1600 |
| Sample Flow Rate (μL/min) | ~250 |
| <sup>m/z</sup>Analyte | <sup>103</sup>Rh |
| <sup>m/z</sup>Internal Standard | <sup>101</sup>Ru |
| DRC Gas | No gas |
| DRC Gas Flow Rate (mL/min) | -/- |
| RPq | 0.25 |
| Mode | MS |
| Nebuliser | Meinhard Plus Series quartz |
| Injector | 1.8mm quartz |
| Spray Chamber | Elemental Scientific cyclonic type |
| Autosampler | Elemental Scientific prepFAST M5 |

### Mass Cytometry (CyTOF)

Mass Cytometry was used in this study to validate metallomic data obtained by SC-ICP-MS from the metal intercalation experiments on Jurkat cells (as described above). In an analogous approach to the SC-ICP-MS analyses, the prepared cell subsets allocated for Mass Cytometry measurements were similarly rinsed three times by centrifugation with CAS+ to fully removal any surface-retained metals and to transfer into the instrument carrier media. All measurements by this technique were conducted using a Standard Biotools Helios CyTOF, which employed the use of its standard single cell suspension pneumatic sample introduction system. The mass cytometer was tuned and its performance confirmed using EQ^TM^ Four Element Calibration Beads. The cells were diluted to 10^6^ cells/mL in CAS+ with 10% EQ^TM^ Four Element Calibration Beads. The .fcs files were acquired and then processed, including normalisation in CyTOF Software v.7 (Standard Biotools).

### Flow Cytometry

Flow Cytometry analysis was incorporated into this study to validate the iron metallomic data obtained by SC-ICP-MS. Cells were transferred to 96 well round bottom plates and washed with PBS. Cells were stained with a cocktail of antibodies and the Zombie NIR fixable viability kit (1:1000, Biolegend, 423105) prepared in PBS for 20 minutes on ice. Cells were subsequently fixed with 2% paraformaldehyde (Pierce, 28906) for 20 minutes on ice, washed and resuspended in PBS. Cells were analysed on either a BD Biosciences LSR Fortessa^TM^ X50 instrument or a Attune NxT flow cytometer (Thermofisher Scientific). Data was analysed using FlowJo (BD biosciences)..

### Statistical Analysis

All individual single cell metallomic mass data was calculated using the single cell module within PerkinElmer’s Syngistix^TM^ ICP-MS software.

Complimentary geometric means and *p* values were calculated in Excel, where the latter was determined using the ‘Regression’ data analysis toolpack. Additionally, other linear regression determinations, such as linear fitting equations and r^2^ values were calculated in R, utilising the dplyr and ggplot2 libraries.

## Acknowledgments and Funding

PH thanks Human Iron Research in Oxford (HIRO), in addition to financial support from PerkinElmer for the funding towards his DPhil. The authors would like to thank Elemental Scientific Inc. for their technical support in commencing analysis with the single cell sample introduction system.

## Author contributions

Conceptualization: HD, JW, MT

Methodology: PH, DP, MT, MM, GP

Investigation: PH, MT, MM, HC, GP

Visualization: PH, GP

Supervision: JW, HD, DP

Writing—original draft: PH, JW

Writing—review & editing: PH, RH, DP, GP, MT, MM, HD

## Competing interests

The authors declare that they have no competing interests.

## Data and materials availability

Tabulated data accompanying the intercalation and murine T-cell experiments are included in the supplementary data at the end of this manuscript. Please contact the corresponding author for further information about the data presented in this manuscript.

## Supplementary Text

### Background correction of SC-ICP-MS data

Single cell inductively coupled plasma mass spectrometry (SC-ICP-MS) rapidly and precisely analyses individual cells from entire cell populations for their intracellular metal contents, thanks to its highly sensitive and fast measurement capabilities. However, SC-ICP-MS data is required to be collected successively, rather than averaged from numerous scans (as per bulk approaches), where corrections to backgrounds (which may exhibit fluctuations) is essential. Dissolved metals found within the extracellular media at levels above the limit of detection will be presented in the time-series data record as a constant signal that is elevated above baseline (*25*). If this signal is stable, systematic 3σ + mean background corrections maybe all that is required to reduce the data to exclusively reveal only the single cell results - as conducted in the Single Cell module of the PerkinElmer Syngistix software. However, other irregular background entities found within the SC-ICP-MS data are much trickier to resolve. An example includes peaks in the data recording fragments of lysed cells and/or those cells that are found below detection. In our study we present the ability to discriminate between such entities and whole cells in the SC-ICP-MS data by virtue of their transit time through the mass spectrometer. Through our rapid scanning procedure, where measurement frequencies are conducted at time-scales that are significantly shorter than the cell events, we are distinctly able to resolve differences between metallomic event durations (and thus whole cells versus cell fragments/ cells below detection). In the experiments that we conducted in this study, our threshold was selected by the lowest peak amplitude that was able to sustain a whole cell event. We found that whole cells above the threshold were presented with well-constrained cell event duration times of ∼0.5ms - when no cell gas is used, to ∼1ms – when the cell is pressurised with NH3. This contrasts those events below the threshold (containing events from cell fragments from lysed cells and/or cell events below detection), which instead transited through the mass spectrometer at much faster time-scales. An example is presented in Fig. S1, where the “non-cell” event in these measurements for iron exhibits a duration time that is more than 30% shorter than the example “cell event”.

Verification towards the accuracy of the background corrections that we applied are highlighted from the transport efficiency data reported from the murine T-cell experiment, where distinct similarities in such values between those collected from calcium to those from iron were made (overall averages being 12.2% versus 12.7% respectively) (see Table S2).

Moreover, as these values are also similar to those transport efficiencies reported from related studies (e.g. *15, 26, 27*), this provides further validation of the method of background correction that we applied.

**Fig. S1.**
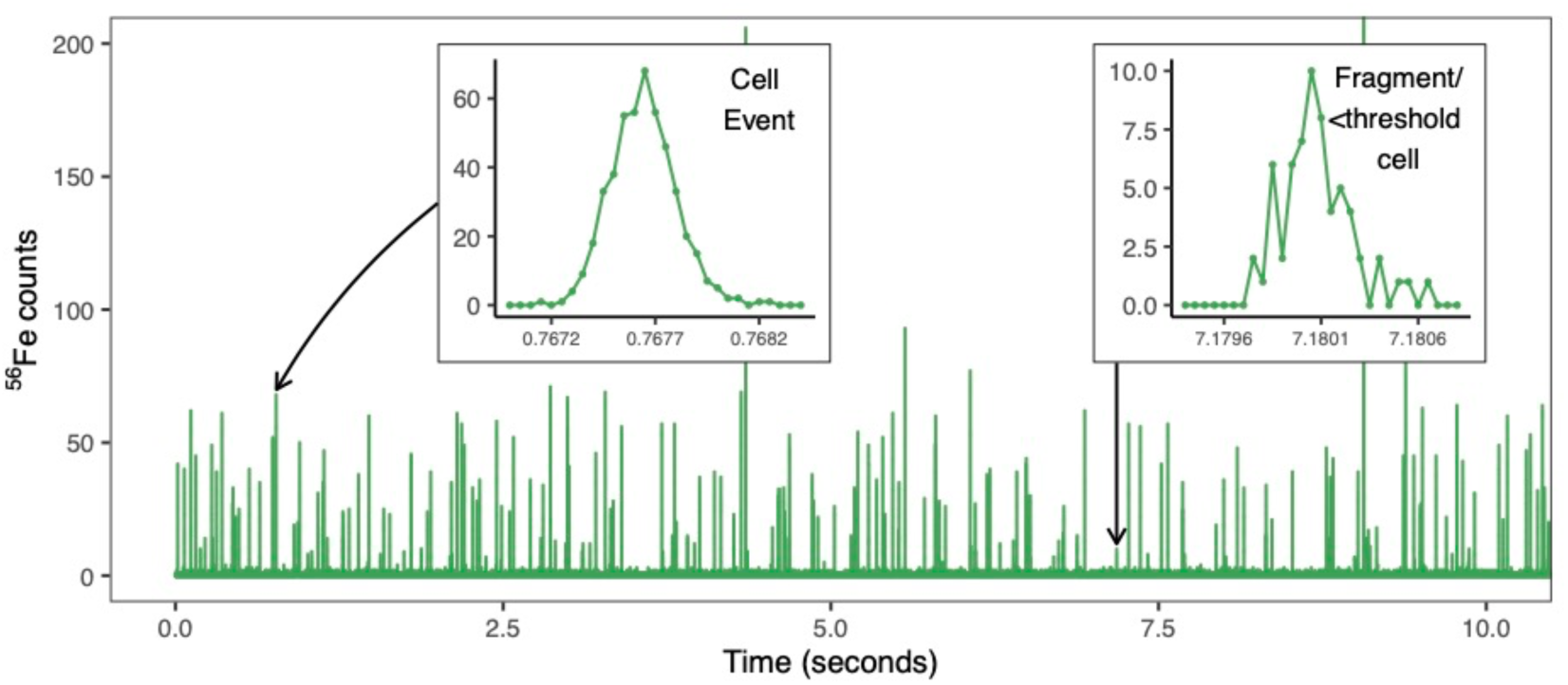
Time-resolved single cell mass spectra measured by SC-ICP-MS from murine T-cells. Measurement of iron in T-cells taken from mouse 3 and the 0.005mg/mL condition. Each spike represents an individual metallomic event, where their individual areas correlate to the mass of iron. The inset taken at ∼0.76 seconds highlights a cell event, where the metallomic plume transits through the mass spectrometer within ∼1ms. The inset taken at ∼7.18 seconds highlights a fragment from a lysed cell/ cell event below detection. Here the metallomic plume transits through the instrument over a much shorter time-span (∼0.65ms), thus rendering it below the detection threshold.

**Fig. S2.**
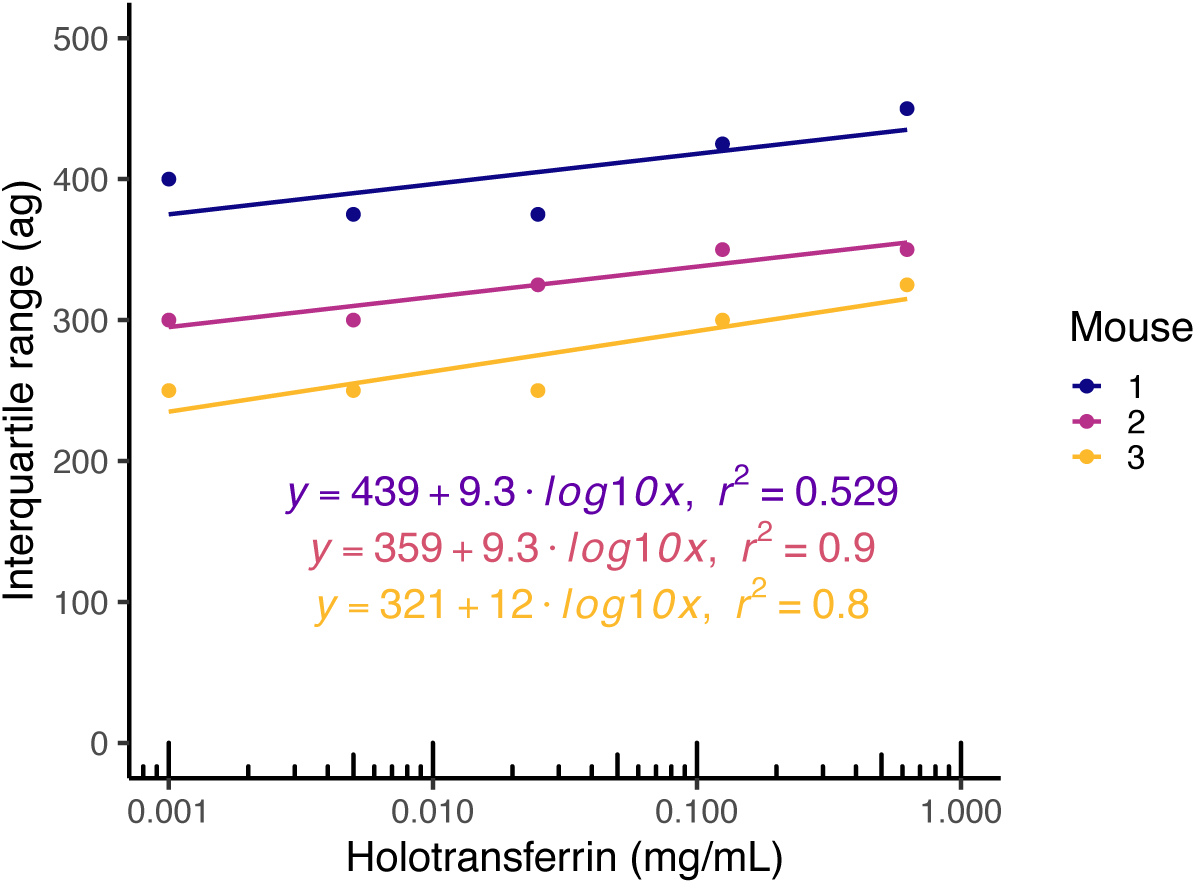
Correlation of inter-quartile ranges against holotransferrin iron-media condition in the murine T-cell experiment. Logarithmic trend lines, associated equations and r^2^ values are also displayed.

**Table S1.** Correlation of inter-quartile ranges against intercalation molarity from the metal-intercalation experiments.

| Intercalation molarity<br>($\mu\text{mol}$) | Metal<br>intercalator | Interquartile<br>range (ag) |
| --- | --- | --- |
| 0.005 | Iridium | 21 |
| 0.01 |  | 7.5 |
| 0.02 |  | 15 |
| 0.03 |  | 24 |
| 0.04 |  | 27 |
| 0.05 |  | 36 |
| 0.05 | Rhodium | 50 |
| 0.1 |  | 48 |
| 0.2 |  | 104 |
| 0.3 |  | 225 |
| 0.4 |  | 320 |
| 0.5 |  | 385 |

**Table S2.**
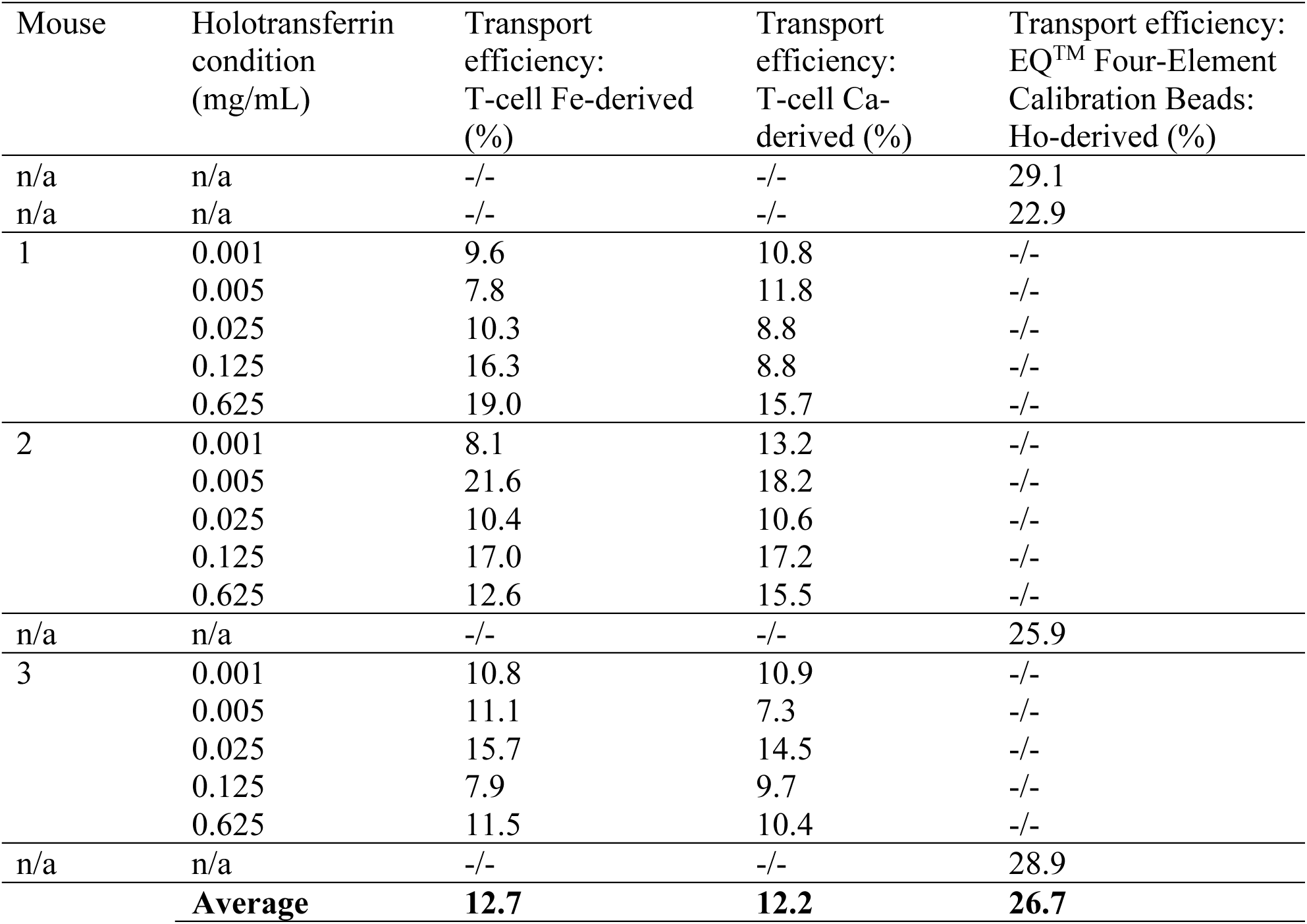
Transport efficiency data from the murine T-cell experiments. All transport efficiency data calculated by the particle frequency method, as described in (*25*). Both iron and calcium-derived transport efficiencies were derived directly from the murine T-cells, and holmium-derived transport efficiencies from EQ^TM^ Four-Element Calibration Beads.

**Table S3.** Live cell counts and total number of cells harvested following the 2-day culturing period for the murine T-cell experiment.

| Mouse | Holotransferrin condition (mg/mL) | Live cell count A (cells/mL) | Live cell count B (cells/mL) | Mean (A and B) (cells/mL) | Total volume (mL) | Total number of cells |
| --- | --- | --- | --- | --- | --- | --- |
| 1 | 0.001 | 1.99E+05 | 2.77E+05 | 2.38E+05 | 7.5 | 1.79E+06 |
|  | 0.005 | 2.67E+05 | 3.40E+05 | 3.04E+05 | 7.5 | 2.28E+06 |
|  | 0.025 | 3.56E+05 | 2.72E+05 | 3.14E+05 | 7.5 | 2.36E+06 |
|  | 0.125 | 2.04E+05 | 2.25E+05 | 2.15E+05 | 7.5 | 1.61E+06 |
|  | 0.625 | 2.30E+05 | 2.83E+05 | 2.57E+05 | 7.5 | 1.92E+06 |
| 2 | 0.001 | 1.88E+05 | 2.77E+05 | 2.33E+05 | 6 | 1.40E+06 |
|  | 0.005 | 3.61E+05 | 3.98E+05 | 3.80E+05 | 6 | 2.28E+06 |
|  | 0.025 | 3.35E+05 | 3.03E+05 | 3.19E+05 | 6 | 1.91E+06 |
|  | 0.125 | 2.20E+05 | 2.67E+05 | 2.44E+05 | 6 | 1.46E+06 |
|  | 0.625 | 2.93E+05 | 2.56E+05 | 2.75E+05 | 7.5 | 2.06E+06 |
| 3 | 0.001 | 4.92E+05 | 5.28E+05 | 5.10E+05 | 7.5 | 3.83E+06 |
|  | 0.005 | 5.91E+05 | 5.02E+05 | 5.47E+05 | 7.5 | 4.10E+06 |
|  | 0.025 | 4.39E+05 | 5.81E+05 | 5.10E+05 | 7.5 | 3.83E+06 |
|  | 0.125 | 4.81E+05 | 4.50E+05 | 4.66E+05 | 7.5 | 3.49E+06 |
|  | 0.625 | 3.92E+05 | 4.08E+05 | 4.00E+05 | 7.5 | 3.00E+06 |

## Notes

### Competing Interest Statement

The authors have declared no competing interest.

